# Polarized Cortical Tension drives Zebrafish Epiboly Movements

**DOI:** 10.1101/032284

**Authors:** Amayra Hernández-Vega, María Marsal, Philippe-Alexandre Pouille, Sebastien Tosi, Julien Colombelli, Tomás Luque, Daniel Navajas, Ignacio Pagonabarraga, Enrique Martín-Blanco

## Abstract

The physical principles underlying the biomechanics of morphogenetic processes are largely unknown. Epiboly is an essential embryonic event in which three distinct tissues coordinate to direct the expansion of the blastoderm. How and where forces are generated during epiboly and how these are globally coupled remains elusive. Here we first develop a method, Hydrodynamic Regression (HR), to infer 3D dynamic pressure fields, mechanical power densities and cortical surface tension profiles within living organisms. HR is based on velocity measurements retrieved from 2D+T microscopy time-lapses and their hydrodynamic modeling. We then applied this method to identify biomechanically active structures during epiboly in the zebrafish and the changes in the distribution of cortex local tension as epiboly progresses. Based on these results, we propose a novel simple physical description for epiboly, where tissue movements are directed by a polarized gradient of cortical tension. We found that this tensional gradient relies on local contractile forces at the cortex, differences in the elastic properties of cortex components and force passive transmission within the incompressible yolk cell. All in all, our work identifies a novel way to physically regulate concerted cellular movements that will be fundamental for the mechanical control of many morphogenetic processes.

## INTRODUCTION

The construction of complex functional structures during morphogenesis results from the unfolding of the developmental program of individual cells. Importantly, morphogenesis does not only rely on a tight regulation of gene expression and cell communication but also depends on fundamental physical principles ^1-9^. The stereotyped tissue movements involved and the embryo optical accessibility makes zebrafish epiboly ideal for exploring mechanical coordination during morphogenesis ^10-12^.

Epiboly is a conserved early embryonic morphogenetic movement that involves a striking physical reorganization characterized by the thinning and spreading of cell layers. Epiboly initiates when the Enveloping Layer (EVL), an external epithelial sheet, the Deep Cells – DCs, a spherical cap of blastomers centered on the animal pole of the embryo over the yolk cell and the External Yolk Syncitial Layer (E-YSL), a specialized region at the yolk surface positioned ahead at the EVL, undergo cortical vegetalward movements. These are associated to the shortening of the rest of the yolk surface (Yolk Cytoplasmic Layer - YCL) and the doming up of the internal yolk. Epiboly ends by the closure of the blastoderm at the vegetal pole covering the yolk cell ^13-15^ **(Fig. 1a)**.

**Figure 1.**
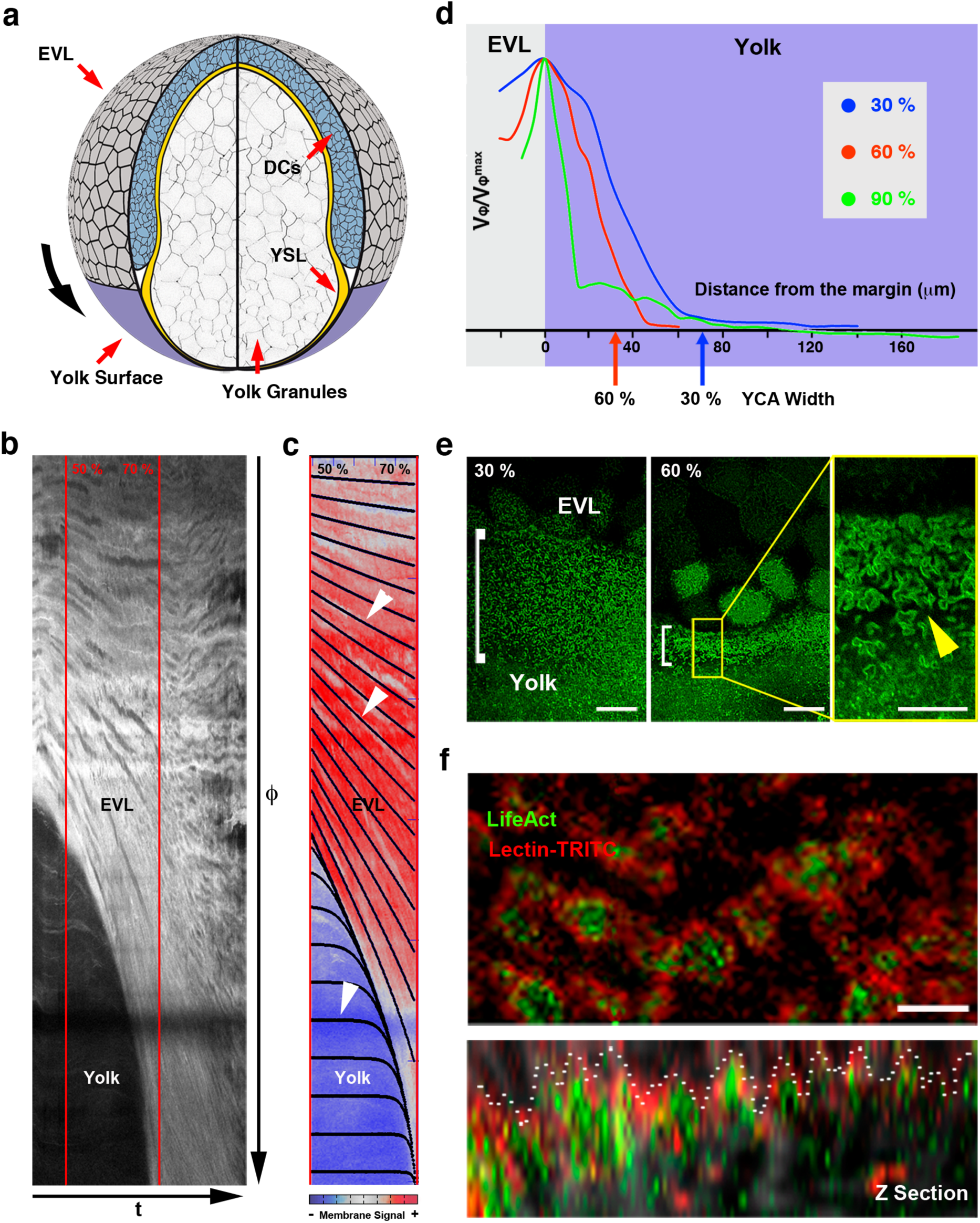
Epiboly kinematics. **(a)** Representation of a zebrafish embryo at 70 % epiboly. Animal (A) and vegetal (V) poles are indicated. The enveloping layer (EVL) is a single-cell layer (dark grey) that stands at the animal cap surface surrounding the inner deep cells (DCs) (blue). It is attached to the yolk cell membrane (purple). The YSL and Yolk Cytoplasmic Layer (YCL) at the periphery and yolk granules fill the yolk cell. **(b)** Kinematics of the embryo surface. A kymograph of the embryo surface (φ axis) from a Tg (β*-actin:m- GFP*) embryo shows the continuous displacement of the EVL / yolk margin, which accelerates after 50% epiboly, while the yolk cell membrane does not show any shift (see **Supplementary Video 1**). **(c)** Theoretical kymograph from 50% to 70% epiboly superimposed, between the red lines, on the experimental kymograph **(b)**. Membrane signal intensity is color-coded (blue to red). White arrowheads point to flow lines (black). **(d)** AV velocities, normalized to EVL / yolk margin velocity, were measured by PIV of time-lapse recordings of the surface of lectin-soaked embryos. The plots correspond to three different epiboly stages (30 %, blue; 60 %, red; and 90 %, green). Blue and red arrows point to the width of the E-YSL at 30 and 60 % epiboly respectively (from **Supplementary Fig. 1**). (e) The yolk cell membrane is highly convoluted (white brackets) ahead of the EVL leading cells and condenses over time. Tg (β*-actin:m-GFP)* embryo at 30 % (left) and 60 % (right) epiboly. Scale bar 25 μm. Inset shows a high magnification of the wrinkled area (arrowhead). Scale bar 10 μm. **(f)** Actin accumulates within wrinkles at the E-YSL. The yolk cell membrane of LifeAct- GFP injected embryos (actin - green) was stained with lectin-TRITC (red) (top). Orthogonal projection of the wrinkled area (bottom). Wrinkles are outlined with a white dotted line. Scale bar 2 μm.

Biomechanical studies, however, experience a major limitation, that is the difficulty to directly measure forces *in vivo* and several methods have been developed to measure the mechanical properties of living cells. So far, however, they cannot be readily applied to whole developing organisms. Exceptions are the use of laser microsurgery ^16^ and the use of genetically engineered mechanical biosensors ^17^. Alas, such techniques are intrusive, fall short of quantitative characterization (e.g. laser cuts cannot distinguish active and passive responses ^18^) and cannot provide a global biomechanical representation of morphogenesis.

We have developed a novel non-intrusive method that we refer as Hydrodynamic Regression (HR), which can accurately and globally estimate the spatio-temporal profile of mechanical power and cortical tension in living organisms. It relies in considering that in deforming tissues during morphogenesis forces are balanced and stresses are continuous at the cortex / fluid interface. This physical principle (boundary condition) has been shown to apply at the single-cell embryo level ^19^ and on multicellular epithelia in gastrulating embryos ^20^. HR is mesoscopic and, although it does not infer the mechanical properties of single cells, it identifies the regions where dynamic stress and mechanical power densities concentrate during tissue rearrangements. Importantly, it does not demand restrictive assumptions either on the mechanical nature of the cortex under stretch or on the physical properties of the embedding fluids. Thus, it can be applied to any morphogenetic process providing two conditions are met: 1) The geometry / shape of the tissues involved can be extracted from images and modeled and 2) The flows around the cortex occur at low Reynolds number, where inertia is negligible, which is the case for most biological processes ^19, 20^.

We applied HR to study epiboly global biomechanics and estimated the topography and temporal evolution of mechanical power density and surface stresses of the embryo. These inferred parameters were validated and confirmed by laser microsurgery, microrheology and Atomic Force Microscopy (AFM). In first place, we concluded that HR is suitable to explore the biomechanics of morphogenetic processes driven by the reorganization of cells or sheets of cells of finite width surrounded by fluids. In countless models, this may most probably be the case. Secondly, the HR analyses and *in silico* modeling let us suggest that epiboly mechanics and kinematics could be simply explained as a result of the integration of the contractile activity of the E-YSL and the differential elastic properties of the EVL and the yolk surface. A tension imbalance will build up across the opposite sides of the E-YSL during epiboly, enabling it to proceed (at least from 50 % onwards). These novel predictions were tested *in silico* and experimentally confirmed by interfering in Misshapen expression (a regulator of myosin contractility active during epiboly progression ^21^) and reducing E-YSL contractility. Our new biomechanical model, in contrast to previously suggested models ^12^, accounts for all known features of epiboly kinematics.

## RESULTS

### Global kinematics of Zebrafish Epiboly

In zebrafish, the EVL, DCs and E-YSL undergo a cortical vegetalward movement throughout epiboly leading to embryo closure (see **Fig. 1a**). Experimental kymographs of the embryo surface along the animal to vegetal axis (φ axis) were extracted from 2D time-lapse meridional sections, perpendicular to the embryos dorso-ventral axis (**Fig. 1b** and **Supplementary Video 1**). These kymographs showed a sustained displacement of the EVL / yolk margin position φ*, whose speed v* continuously increased and that after crossing the equator (v_0_) could be fitted in theoretical kymographs simulations to the function v*_ϕ_* = v_0_ / sin *ϕ* * **(Fig. 1c)**. Theoretical kymographs were built from known kinematic data and designed to maintain the elastic equilibrium state (see **Supplementary Note 1**). Experimental kymographs also showed that from the EVL / yolk margin towards the vegetal pole, the local surface speed decreased exponentially **(Fig. 1d)**. This decay delineates two distinct domains, a proximal one ahead of the EVL, roughly equivalent to the E-YSL, which undergoes progressive constriction (see **Supplementary Fig. 1**) and a distal mostly static area. The proximal E-YSL constitutes a morphologically distinguishable region showing a highly convoluted yolk cell membrane ^22, 23^ **(Fig. 1e)**. In the E-YSL, actin was conscripted within and beneath wrinkles **(Fig. 1f)** underlined by nestled mitochondria (see also ^24^) (**Supplementary Fig. 2**). Importantly, we observed that the width of the E-YSL (*δ ϕ*) and the velocity of the EVL / yolk margin v* were inversely paired (v* ~ 1 / *δ ϕ*) at all times. While this pairing is just qualitative up to reaching the equator (v_0_), it does quantitatively match the theoretical kymographs after 50 % **(Fig. 1c)**.

**Figure 2.**
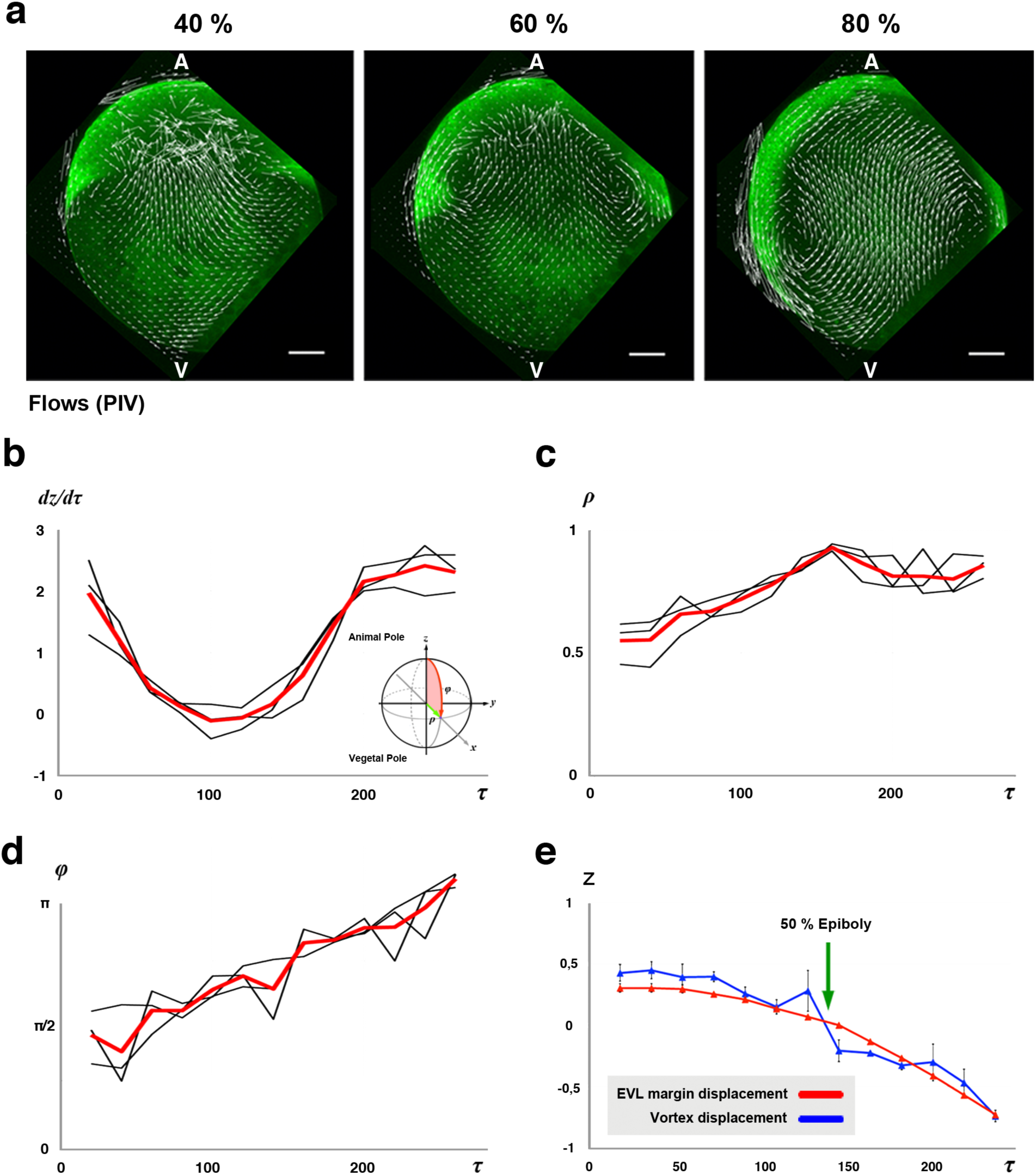
Yolk Flows. **(a)** Yolk granules flow in stereotyped patterns as epiboly progresses. PIV of time-lapse snapshots imaged by two-photon microscopy of a Tg (β*-actin:m-GFP)* embryo (from **Supplementary Video 2**). Upon epiboly onset, yolk granules flow towards the animal pole displaying two internal vortices near the E-YSL. At 50 %, the movement of the yolk granules halts, but resumes afterwards until the end of epiboly. Animal (A) and vegetal (V) poles are indicated. Scale bar 100 μm. **(b)** Internal yolk content speed (μm/min) at the center of the embryo, over time (min). Inset: Spherical coordinates. The main axis of the sphere is represented by z, from the animal to the vegetal pole. The polar angle is represented by φ and the radial coordinate is denoted ρ. (c) Vortex midpoints distance from the center (ρ), over time (min). **(d)** Vortex midpoints position in relation to the φ axis, over time (min). Black lines represent individual measurements. Red lines represent averages. **(e)** Correlation between vortex displacement (blue, average of n=3 embryos) vs. EVL margin displacement (red, average of n = 4 embryos) along the main axis (z). Standard errors are displayed. Arrow points to 50 % epiboly.

Further, we observed that coupled to EVL, DCs and E-YSL epiboly, internal yolk granules also undergo stereotyped motions. These granules sustain a spinning symmetrical movement around a transverse annular axis creating a torus of vortices, running from the vegetal to the animal pole at the center and from the animal to the vegetal pole at the periphery (**Fig. 2a** and **Supplementary Video 2**). Vortices centroids move gradually towards the cortex and towards the vegetal pole as epiboly progresses (**Fig. 2b-d**). Importantly, they parallel the displacement of the edge of the EVL along the *ϕ* axis, which after 50 % epiboly progresses at a constant velocity of 200 μm/hour towards the vegetal pole (z axis) **(Fig. 2e)**.

In summary, the EVL leading edge movements, the shrinking and displacement of the E-YSL and the stereotyped vortices generated by the movements of the yolk granules are remarkably coordinated and seem mechanically coupled during epiboly.

### Development of a non-invasive method, Hydrodynamic Regression, to determine mechanical power densities and stress patterns

Although appears to be clear that one of the main power sources for epiboly is the progressively confined actomyosin cortical belt generated at the E-YSL cortex ^21, 23^, how tissue behaviors mechanical coordination is achieved remains unexplored. To survey this mechanical coupling we developed a mesoscopic method of general applicability (Hydrodynamic Regression) able to extract dynamically mechanical power density maps and embryo cortex stress patterns.

#### Method Rationale

Cortices in developing systems, as their width is much smaller than the organism size, can be mesoscopically described as continuous quasi-2D structures under tension embedded in viscous fluids. As such, when a cortex expands or contracts, its mechanical dynamics (surface tension differences) can be derived from the shear stress values at its interface. Shear strain (deformation) measurements near the cortex surface for most morphogenetic processes are, however, difficult to obtain. Thus, we reasoned that we could indirectly infer the local shear stresses generated by cortical deformations by analyzing the flows in the surrounding viscous media **(Fig. 3a)**. More precisely, any variation in the tangential stresses at the sides of the cortex, δ**σ**_tn_ will be quantitatively proportional to the 2D out-of-equilibrium local cortical tension τ (unbalanced cortical stresses). Tangential stresses, due to shear viscosity and the stress balance at the cortex, will be equal to the shear stress of the adjacent fluid. Therefore, by analyzing the velocity fields of embedding fluids and by determining the fluid shear stresses at the cortex interface, it should be in principle possible to infer the quantitative profile of τ up to an additive constant (see below).

**Figure 3.**
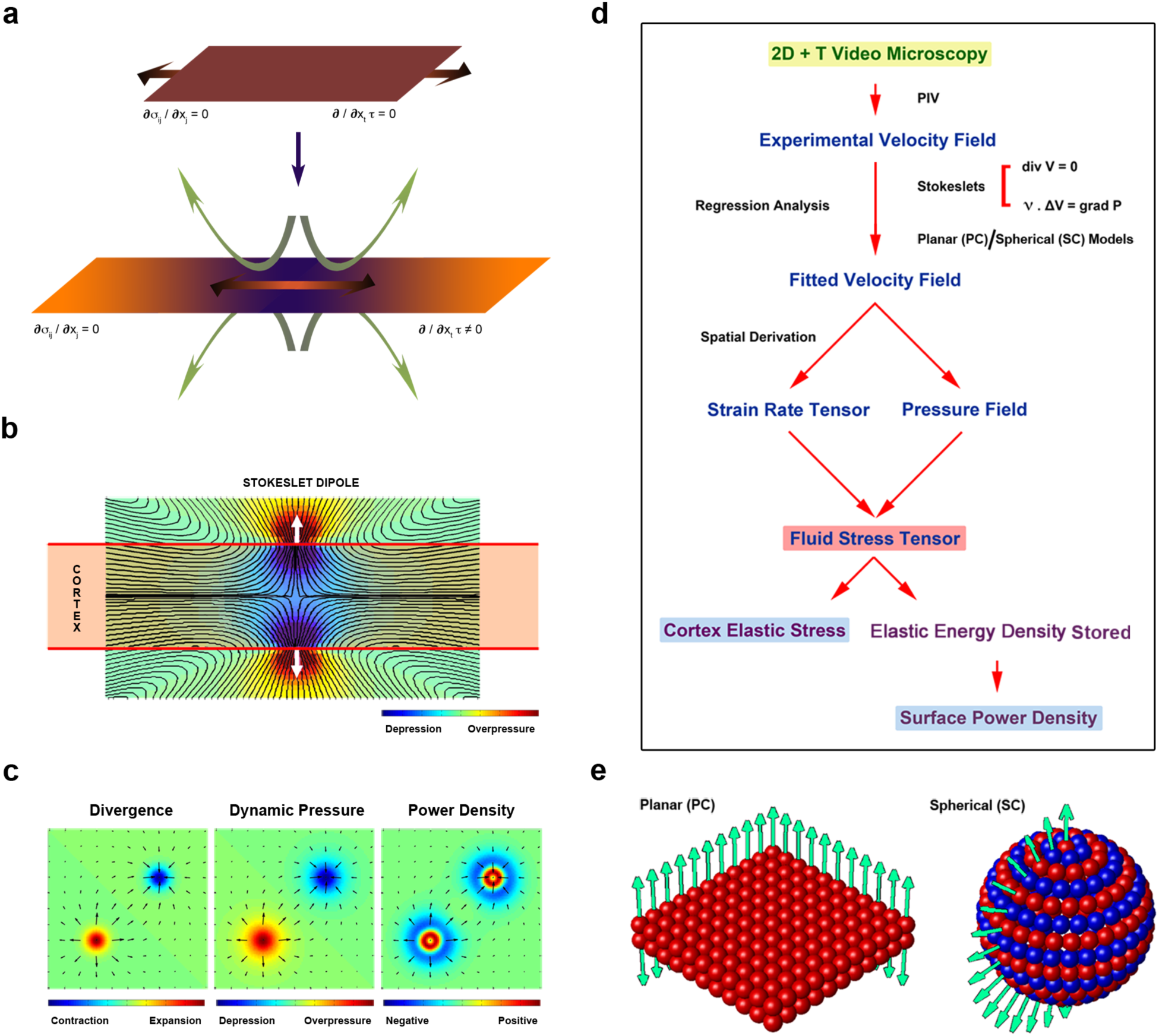
Hydrodynamics Regression. **(a)** Any cortical surface of living organisms could be considered a structure under tension embedded in a viscous liquid, where the stresses in the three dimensions at each point are locally balanced *∂***σ**_ij_ / ∂x_j_ = 0 and the cortex tension τ is at equilibrium ∂ / ∂x_t_ τ = 0 (top). Any disequilibrium in this tension (∂ / ∂x_t_ τ ≠ 0), e.g. due to the stretching of the cortex surface (graded double-headed arrow), will result in a discontinuity of the shear stress of the surrounding viscous fluid and the induction of flows (green arrows) (bottom). Thus, the measurement of the shear flow of the viscous fluid on both sides of a cortical surface lets inferring the quantitative profile of the surface tension along the cortex, regardless of its elastic properties. **(b)** Pipeline describing the different steps involved in HR. From time-lapse 2D videos (green shadowed in yellow) a set of parameters for the fluid phase (blue) are sequentially acquired through PIV and regression analysis, in particular the fluid stress (shadowed in pink). From these, the relative elastic stress and mechanical power density (purple shadowed in pale blue) of the cortex are extracted for different geometrical configurations **(c)** Representation of an Stokeslet pair, at the limits of a perpendicular section of a planar hydrodynamic active cortex (red shadow), pointing in opposite directions (white arrows) and their associated flow lines (black) and dynamic pressure fields (color-coded). **(d)** Divergence fields of Stokeslet dipoles of opposite magnitudes corresponding to cortical expansion or contraction (left). These dipoles carry an associated dynamic pressure (middle) and power density (right). Mechanical power density is positive at the central Stokeslet pair that is surrounded by a cortical ring that resists its deformation. **(e)** Representation of the planar cortex (PC) (left) and the spherical cortex (SC) (right) models where the elementary solutions (dots) pointing in opposite directions (pale green arrows) are evenly distributed on a square grid (PC) or symmetrically distributed around the main cylindrical axis of a sphere (N_c_ = 16) (SC). For the PC model only the external components are shown in one row and one column. For the SC model only the external components are shown in one column.

We employed regression analysis (HR) to analytically fit modeled velocity fields in and outside the cortex to experimental velocity fields. HR let to retrieve the overall spatiotemporal distribution of τ and the corresponding dynamic pressure distribution in the fluid and at the fluid-cortex interface. Theoretical considerations and derivations are fully detailed in **Supplementary Note 2**.

#### Method implementation

We first determined the velocity fields of the media surrounding the tissue/cortex. To do so, we employed bright field transmission or two-photon confocal microscopies and estimated tissue kinematics by Particle Image Velocimetry (PIV) ^25^. PIV allows extracting local displacements (velocity fields) between consecutive movie frames. Second, we defined the geometry of the cortex and modeled the surrounding fluid motions by distributing a given number, N_e_, of force dipoles (Stokeslet pairs) of variable strength β_e_ perpendicular to the cortex and at a distance δ away from each other **(Fig. 3b)**. These force dipoles account for the cortical expansions or contractions associated to the motion of nearby fluids and let to model the 3D fluid velocity field and its corresponding dynamic pressure as a linear combination of the fluid flows generated by each pair. This modeling assumed that in most developmental processes fluids at small strain rates behave as incompressible Newtonian fluids at low Reynolds number (REFS, ^26^). Fluid dynamics can then be described by a linear continuity equation, ensuring volume conservation, and the linear Stokes equation

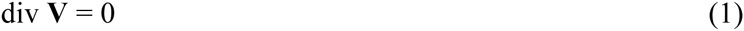

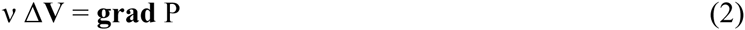
with *v* being the shear viscosity, **V** the velocity field, Δ the Laplace operator, **grad** the gradient and P the dynamic pressure.

Third, we used regression analysis to fit the analytical theoretical model to the experimental velocity field. At each time point a Monte Carlo scheme was used to determine the values of the parameters β_e_ that minimize the mean square error (MSE) between the experimental and theoretical fields. The corresponding dynamic pressure was determined from these same values.

Last, considering the fluid strain rates and the dynamic pressure values, we inferred the fluid shear stress, the local values of *τ*, the average surface tension and the cortical mechanical power density Π, *i.e*. the rate of mechanical energy produced by the unbalanced cortical tension per unit of time **(Fig. 3c)**.

Performing HR over time gives access to the spatio-temporal evolution of both, cortical stresses and mechanical power density maps. The pipeline used to infer the overall mechanics of a cortex and its surrounding fluids by fitting HR models to experimental flows velocity measurements is shown in **Fig. 3d**. The workflow and protocols involved are thoroughly described in **Supplementary Note 2**. A flow simulation Software suite with step-by-step details on the procedure to use the code is provided in **Supplementary Note 3**.

HR can, in principle, be applied to different scenarios, materials and geometries. We explored this versatility by analyzing as benchmarks diverse 2D and 3D active structures with known or easily predictable mechanical behaviors. We examined the reaction of the external yolk membrane / actomyosin cortex of the zebrafish embryo to laser injury by employing planar cortex (PC) models developed by positioning stokeslet pairs on a 2D square grid. Further, spherical cortex (SC) models built by evenly distributing stokeslet pairs around one main axis on a spherical shell with cylindrical symmetry were employed to evaluate a spherical fluid droplet sedimenting in an immiscible liquid (**Fig. 3e** and **Supplementary Note 4**). In both cases, HR appears to faithfully infer dynamic pressure fields, mechanical power densities and surface tension maps from experimental velocity fields (see **Supplementary Fig. 3** and **4** and **Supplementary Video 3**).

### Epiboly Biomechanics

We performed a thorough biomechanical study to define the active elements in and around the E-YSL guiding and/or contributing to epiboly progression.

#### Yolk mechanical properties

We first assessed the mechanical properties of the yolk by studying its rheological behavior recording the thermal fluctuations of microinjected fluorescent nanoparticles ^27^. Two hours after microinjection, images of yolk-injected nanoparticles were obtained and the position of the centroids, the trajectories and the two-dimensional mean square displacement (MSD) of each particle were computed. We found that the MSD exhibited a proportional dependence on time lag for the yolk consistent with Newtonian fluid behavior and a viscosity of 129 mPa.s **(Fig. 4a)**. This viscosity is in the same order to that recently determined for the *Drosophila* yolk (286 mPa.s) ^20^. Considering this viscosity, the speed of the yolk flows referred to epiboly progression rates (200 μm/hour), the yolk density (~ 1 g/cm^3^) and the size of the embryo (~ 350 μm of radius), we calculated a Reynolds number for the yolk in the order of 10^-7^ (as in *Drosophila* ^20^), a very low value.

**Figure 4.**
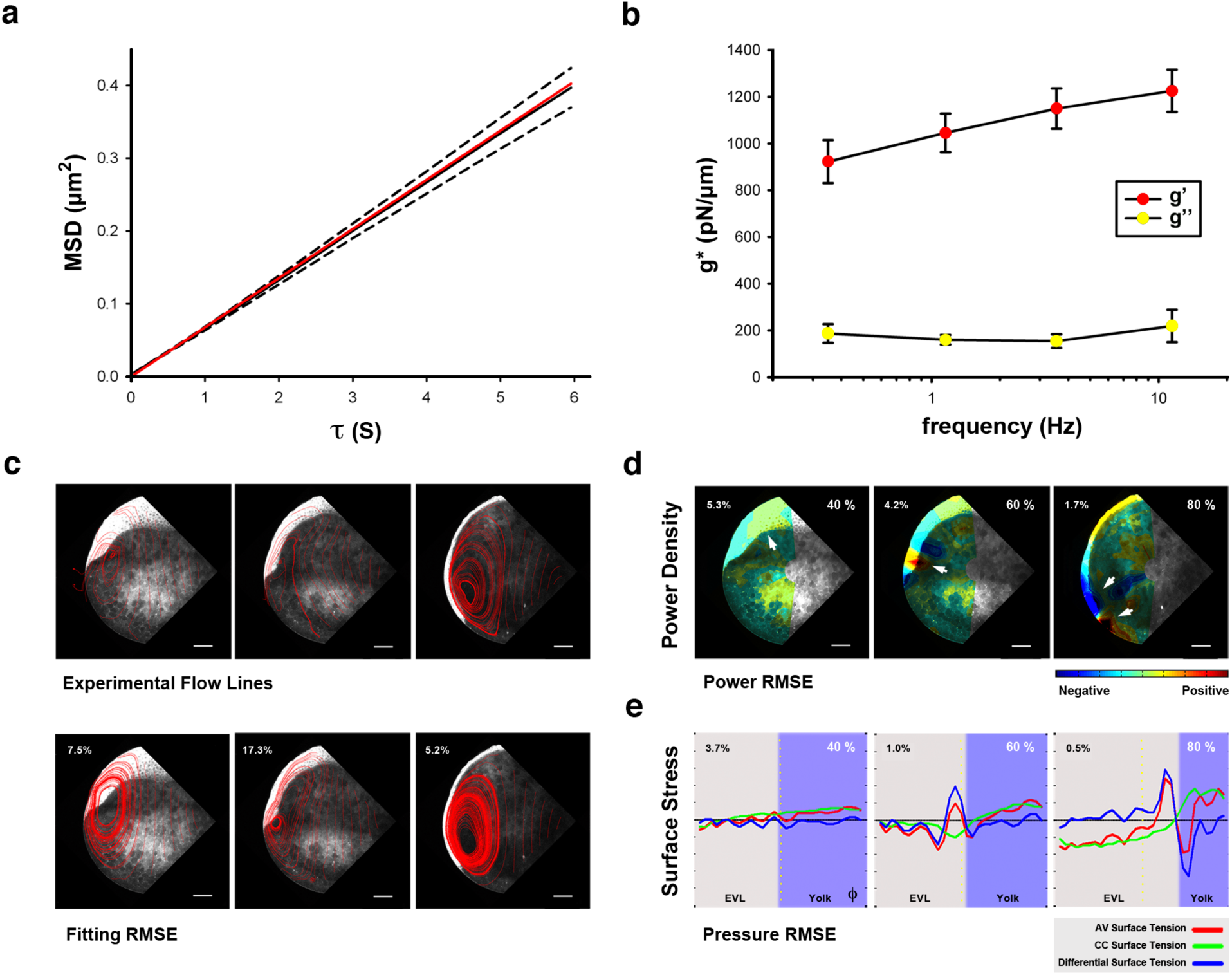
Epiboly biomechanics. **(a)** Yolk microrheology. Mean squared displacement (MSD) of thermal fluctuations of fluorescent nanoparticles embedded in the yolk of zebrafish embryos. Solid and dashed black lines are mean ± SE computed from 99 particles. Red line is the fit of the Stokes-Einstein equation. MSD of nanoparticles embedded in the yolk exhibits a proportional dependence on time lag, τ, consistent with a Newtonian liquid behavior with a viscosity of 129 cP. A linear time dependence of MSD with similar values of yolk viscosity has been recently reported in *Drosophila* ^20^. Interestingly, the viscosity of the zebrafish yolk is two orders of magnitude lower than that of C. *elegans* embryos ^27^ and of the cytoplasm of several cell types ^40^. **(b)** Viscoelasticity of the yolk cortex. Complex modulus (g*) measured by indenting the surface of zebrafish embryos with AFM. Solid symbols are the elastic modulus and open symbols are the viscous modulus, respectively. In all the measured frequencies cortex rheology is dominated by a solid-like behavior with the viscous modulus 5 times lower that the elastic modulus. No significant frequency dependence was found in the elastic (ANOVA, p = 0.123) and viscous modulus (ANOVA, p = 0.719). **(c)** Flow lines from snapshots of the PIV measurements of a two-photon excitation video time-lapse (from **Supplementary Video 4**) of a Tg (β*-actin:m-GFP)* embryo (top), and fitted flow lines using HR (bottom). The relative mean square error (RMSE) of the fitting is displayed as a percentage. Scale bar 100 μm. **(d)** Mechanical power density maps over time obtained by HR (from **Supplementary Video 5**). The RMSE of the power is shown as a percentage. The mechanical work to produce doming (40 % epiboly) arises mainly at the blastoderm (left); at 60 % epiboly (middle) DCs undergo ingression (red) while the E-YSL resists constriction (blue). At 80 % epiboly (right), the active work leading to the active movement of the EVL / yolk margin mainly maps at the E-YSL (red) while the blastoderm undergoes elastic resistance (blue). White arrows point to the main observed events. Scale bar 100 μm. **(e)** AV (red) and CC (green) stresses and their differences (blue) (from **Supplementary Video 6**) along the embryo cortex were plotted as a function of the 9 angle from animal to vegetal (from the regression analysis). The equator - dotted yellow line -, yolk surface - purple shadow - and the RMSE of the dynamic pressure as a percentage for each time point are displayed (40 %; 65 %; 80 % epiboly). The CC stress steeps up from animal to vegetal and the AV shows a more complex profile. Their difference changes senses around the EVL / yolk margin.

We then performed a direct quantification of the yolk cortex viscoelastic properties. Atomic Force Microscopy (AFM) was employed to acquire local surface tension measurements on living whole embryos (see **Supplementary Note 5**). The viscoelasticity of the cortex was assessed applying low amplitude (100 nm) multifrequency oscillations composed of sinusoidal waves of different frequencies. We found that in all measured frequencies (0.35 Hz – 11.45 Hz), cortex rheology was dominated by an elastic-like behavior with the viscous modulus 5 times lower than the elastic modulus **(Fig. 4b)**. Moreover, no significant frequency dependence was found for any of these moduli in the suitable experimental time window. As the ratio between the viscous and elastic moduli is expected to drop with decreasing frequency, our rheological data indicated that at a time scale of minutes the cortex, with an elastic modulus of 100 Pa, behaves as a very soft solid in the range of the softest biological materials ^28, 29^.

Last, from the measured relaxation times determined by AFM we estimated the effective cortex viscosity at long-time scales (fluid-like response) in the order of *η_c_* ~ 10 Pa.s. This implies that the ratio between the typical traction in the cortex at long times and the stresses associated to the measured flows of the fluid is just about *η_c_w / nR* ~ 0.1, where *w* stands for the cortex width, R the characteristic embryo size and *η* the yolk viscosity.

In conclusion, the yolk content can be considered a viscous Newtonian fluid with low Reynolds number and the yolk surface behaves at the time scale of epiboly movements as an extremely soft solid. Importantly, the magnitudes of the cortical and yolk fluid tractions are truly comparable.

#### Epiboly mechanical power sources

We applied HR to uncover the spatiotemporal pattern of mechanical power density during epiboly. To do so, experimental 2D velocity fields from time-lapse videos of embryos meridional sections were determined by PIV employing yolk granules movements as a reference. Then, simulated 3D velocity fields were generated from a SC model with Stokeslet pairs distributed on five concentrical spherical shells adjusted to the epiboly motion symmetry and the embryo cortex. These theoretical fields were eventually fitted by regression analysis to the experimental velocity fields (**Fig. 4c** and **Supplementary Video 4**).

We found that at the epiboly onset, the active power mainly maps to the blastoderm. Later, a distinct high power domain arises under the cortex at the position and time of the ingression of DCs between 50 and 60 % epiboly. Last, once the EVL front crosses the equator, the largest mechanical power localizes to the actomyosin-rich E-YSL, while the EVL adjacent cells display negative power densities opposing deformation. The EVL cells and DCs near the animal pole remain a weak power source (**Fig. 4d** and **Supplementary Video 5**). At all times, the yolk always displays very low mechanical energy dissipation.

#### Cortical Stress Patterns

HR enabled to analyze the spatio-temporal progression of local surface tension by linearly mapping animal to vegetal (AV) and circumferential (CC) cortical stresses as a function of time (**Fig. 4e** and **Supplementary Video 6**). We found that at the initiation of epiboly both, AV and CC stresses show an even distribution at the surface of the embryo. As epiboly progresses, however, a vegetalward-oriented gradient of CC stress is built up throughout the EVL and the yolk, while the AV stress displays a complex distribution. AV diminishes with time at the animal pole, dramatically rises at the EVL leading front, drops at the E-YSL and increases again towards the vegetal pole. As a result, after 50 % epiboly, the difference between AV and CC stresses resulted in sustained minimum negative values at the E-YSL. In contrast, the stress difference in the adjacent EVL cells was positive and maximum. Summing up, a positive gradient of tension progressively develops at the yolk surface towards the vegetal pole from 50 % up to the end of epiboly. This is more pronounced for the CC component.

The tension pattern predictions were validated by targeted laser microsurgery ^30^ (**Supplementary Figs. 5** and **6** and **Supplementary Note 6**). The experimental disruption in the AV or CC directions of the yolk actomyosin cortex at the E-YSL resulted in immediate recoil with exponentially decaying speed (see also ^12^). This enabled to measure the relative magnitude and directionality of local surface tensions (Supplementary **Fig. 7** and **Supplementary Video 7**). Significantly, we observed that the differences between AV and CC tension at each time point precisely replicated the stress differences estimated by HR **(Fig. 5a)**. Thus, the inferred temporal and directional patterns of tension at the E-YSL were confirmed. Equally, the vegetalward AV tension gradient at the yolk surface predicted by HR was also experimentally confirmed by laser cuts performed in the yolk at different distances from the EVL / yolk margin and at different stages **(Fig. 5b)**.

**Figure 5.**
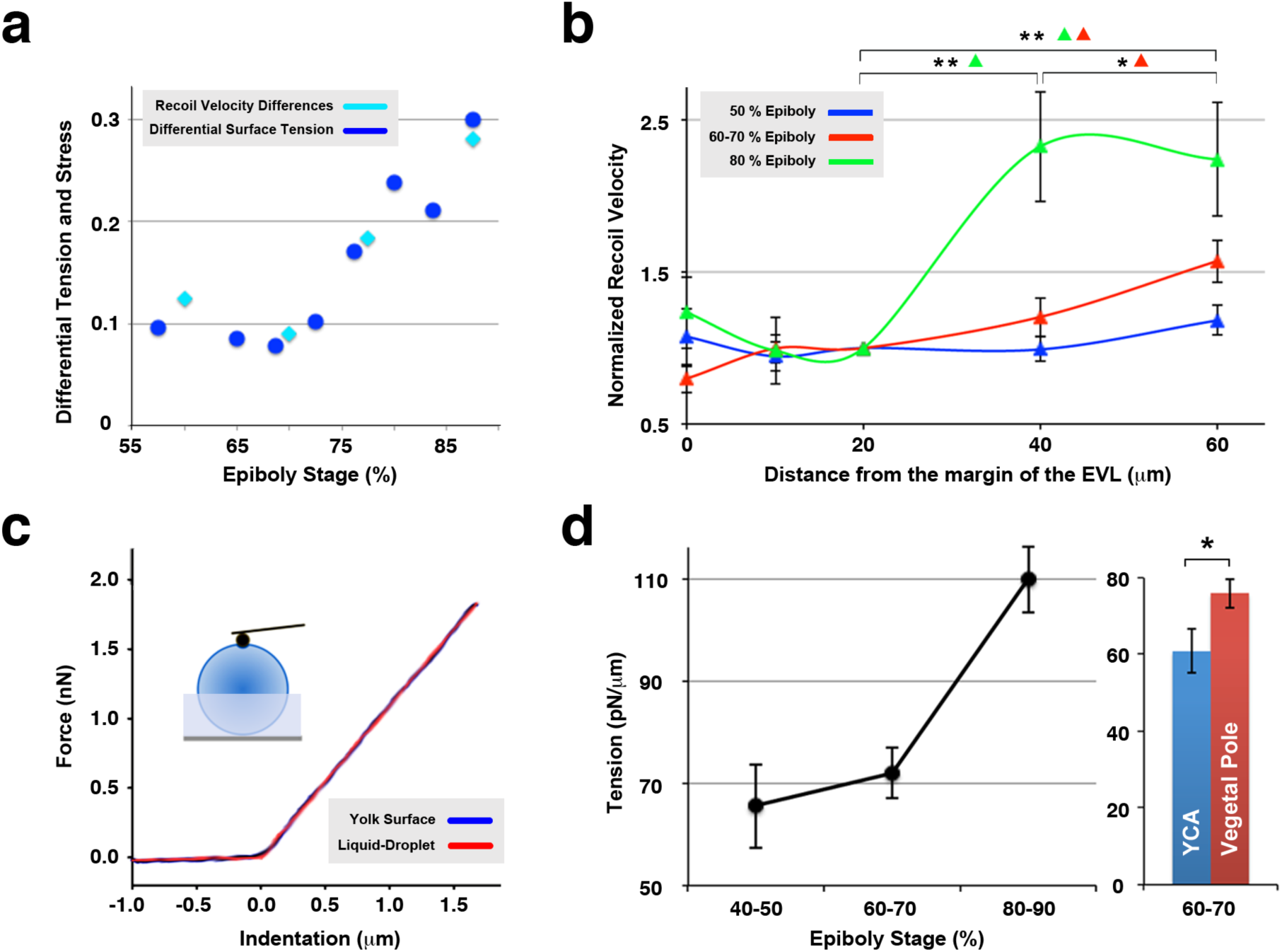
Experimental validation of epiboly tensional pattern. **(a)** Correlation between the differential tension (cyan) represented by the recoil velocities differences (μm/s) between CC and AV laser cuts and their difference (blue) calculated by regression analysis at the E-YSL during epiboly (%). The difference values were fitted applying a constant adjustment ratio at all time points. **(b)** AV tension in the yolk cortex at different distances from the EVL (0, 10, 20, 40 and 60 μm) and at different epiboly stages (50 %, blue; 60–70 %, red; 80 % epiboly, green). Significant changes are highlighted. **(c)** Probing mechanical parameters of living embryos with AFM. Force-indentation curves recorded at cantilever velocity of 10 μm/s (blue) fitted with a "liquid balloon" model (red). Inset: AFM force-indentation description. **(d)** Averaged values of mean surface tension (pN / μm) on the yolk obtained with AFM at different epiboly stages (left), and the significant difference of tension at 60–70 % epiboly (p = 0.0245) between the E-YSL and the vegetal pole (right).

Stress-graded profiles were also confirmed by AFM. We measured force-indentation curves at the yolk surface and found that they showed a perfect match to a "liquid balloon" model. This is a viscoelastic mechanical model representing a viscous liquid surrounded by an elastic cortex under mechanical stress ^31-33^. By fitting this model to the experimental data we calculated absolute values of the cortical tension (Tc) of the embryo yolk membrane in the order of 100 pN / μm **(Fig. 5c)**. This analysis (on two different positions, between 50 and 100 μm ahead of the EVL and around the vegetal pole) supported a significant long-range positive gradient of tension after 50 % epiboly and unequivocally pointed to the presence of tension at the vegetal pole **(Fig. 5d)**. These results are consistent with the short-range gradient observed by laser cuts and in agreement with the HR analysis.

In short, the yolk surface stresses and dynamics predicted by HR were accurately supported by AFM and laser microsurgery.

### Mechanically relevant elements leading epiboly movements

The differences in local surface speeds observed in the kinematic analysis suggest potential stiffness inhomogeneities at the embryo cortex that could be important for the understanding of epiboly mechanics. We surveyed this option by studying the cortex response to 20 μm wide laser cuts performed on the yolk membrane parallel to the EVL and found, after recoil, a consistent displacement of the geometrical center of these cuts towards the vegetal pole **(Fig. 6a)**. These data not just support the presence of a tension gradient but, importantly, highlight the difference of stiffness between the EVL (softer) and the yolk (stiffer) surfaces. This difference correlates with previous analysis of micro-elasticity using Brillouin Scattering Spectroscopy, which also revealed the relative higher stiffness of the yolk versus the blastoderm at early cleavage stages ^34^. Overall, the kinematics indicates that the EVL behaves mesoscopicaly as a homogeneous deformable sheet, that the E-YSL shrinks specially after 50% epiboly, and that the rest of the yolk cell membrane is mostly un-stretchable and kept under an increasing tension.

**Figure 6.**
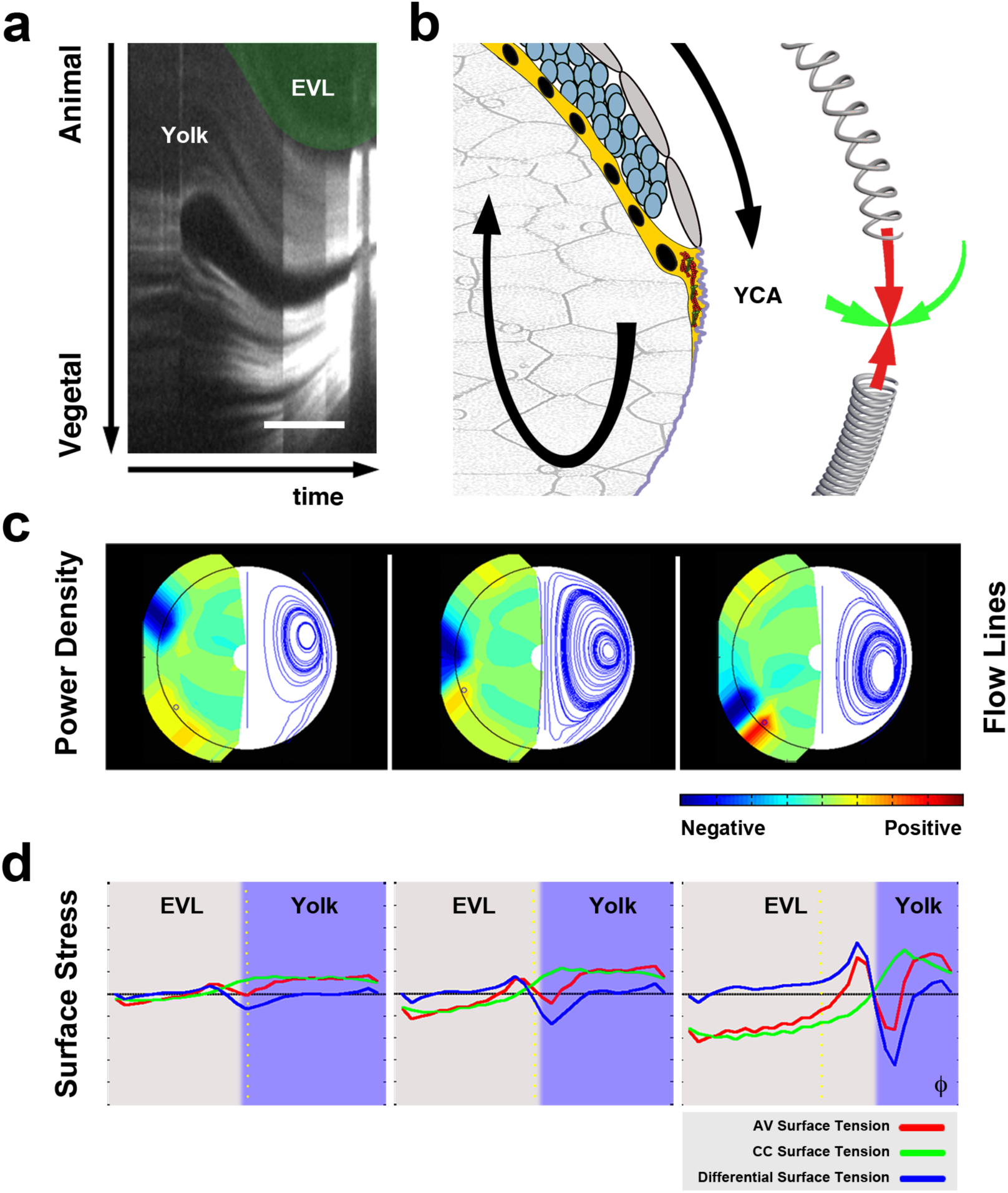
Epiboly is driven by E-YSL contractility and differential surface tensions. **(a)** Kymograph showing the deformation of the cortex along a segment perpendicular to a laser cut in the E-YSL. After a rapid opening, an active contraction displaces the cut centroid vegetalward indicating that the EVL is more easily stretched (elastic) than the vegetal yolk cell membrane/cortex. When the gap is closed, the cut center returns to its initial position and the elastic equilibrium is restored. Scale bar 10 μm. **(b)** Proposed model of epiboly progression. The contractile E-YSL and the imbalance of stiffness between the EVL and the yolk surface account for epiboly progression. The vegetal boundary of the E-YSL is not mechanically free but linked to the tense yolk membrane. The tension originated by constriction of the actomyosin ring (E-YSL) is thus conveyed by the yolk membrane towards the vegetal pole leading to the progression of the less stiff EVL. Passive movements within the yolk arise as a result of force transmission from the surface. **(c)** Dynamic simulation of epiboly based on a single spherical cortex model (SC) considering a gradient of elastic tension from animal to vegetal and a constricting area at the E-YSL (from **Supplementary Video 8** - 40 %; 60 %; 80% epiboly). Color-coded mechanical power density map (positive - red, negative - blue) (left). Flow lines (right). The mechanical power is supplied at the surface and internal vortices and their shift from animal to vegetal are precisely simulated. **(d)** Dynamic simulation of AV and CC stresses (from **Supplementary Video 9** - 40 %; 65 %; 80 % epiboly) represented as in **Fig. 1a**. The AV and CC stresses are equal at the poles, as the epiboly simulation progresses, the CC stress steeps up from animal to vegetal and the difference between CC and AV stresses change sense at the EVL / yolk margin.

Based in these and previous data, we hypothesize that epiboly, at least after 50 %, will be just reliant on the active contraction of the E-YSL and directed towards the vegetal pole as a reaction to the difference of stiffness between the animal (EVL) and vegetal (yolk) domains, which would result in the establishment of a cortical tension gradient **(Fig. 6b)**. As a first assessment of this hypothesis we simulated epiboly *in silico* via HR (SC model). This simulation took into account the experimental cortical kinematics and enforced an exponential decay of the cortical contractility along the φ axis (set in accordance to the observed exponential decay of speed of the EVL / yolk margin) and assumed that both the EVL and the yolk cell membrane behave as materials of different stiffness. We then considered a spherical shell representing the cortical surface of the virtual embryo, set the EVL / yolk margin to its position at the onset of epiboly and imposed *ad hoc* β_e_ coefficients to the Stokeslet pairs to fit a simulated velocity field to the experimental velocity fields. The variation of Stokeslets strength, β_e_, with their position along the dorso ventral axis (imposed in the simulation) accounts for the different stiffness of the EVL and yolk cortex. The exponential decay of contractility is specifically restricted to the E-YSL by imposing positional limitations for this term. For each time step of the simulation, the longitudinal component of the velocity vector v* was used to shift forward the EVL / yolk margin (see **Supplementary Note 7**).

We found that SC model simulations accurately recapitulated, at least from 50 % epiboly onwards, most of the kinematic and mechanical parameters associated to epiboly, the yolk internal vortices, the cortex mechanical power densities and the patterns of surface tension at the cortex (**Figs. 6c** and **d** and **Supplementary Videos 8** and **9**). This mechanical model would predict that interfering with the contractile capabilities of the E-YSL, i.e. by abolishing the progressive reduction of its width, would result in a dramatic reduction in the speed of the advancing front and, in practical terms, in epiboly completion failure.

We found that imposing the experimental kinematic regime of E-YSL shrinkage (with a two step decay) to SC model simulations rendered an specific profile for the speed of the EVL front displacement along the φ axis remarkably accurate, first modest up to reaching the equator and exponential afterwards. Importantly, simulations in which the E-YSL shrinkage was abolished resulted in a completely different regime showing a progressive decline of the front speed from 50 % epiboly onwards **(Fig. 7a)**.

**Figure 7.**
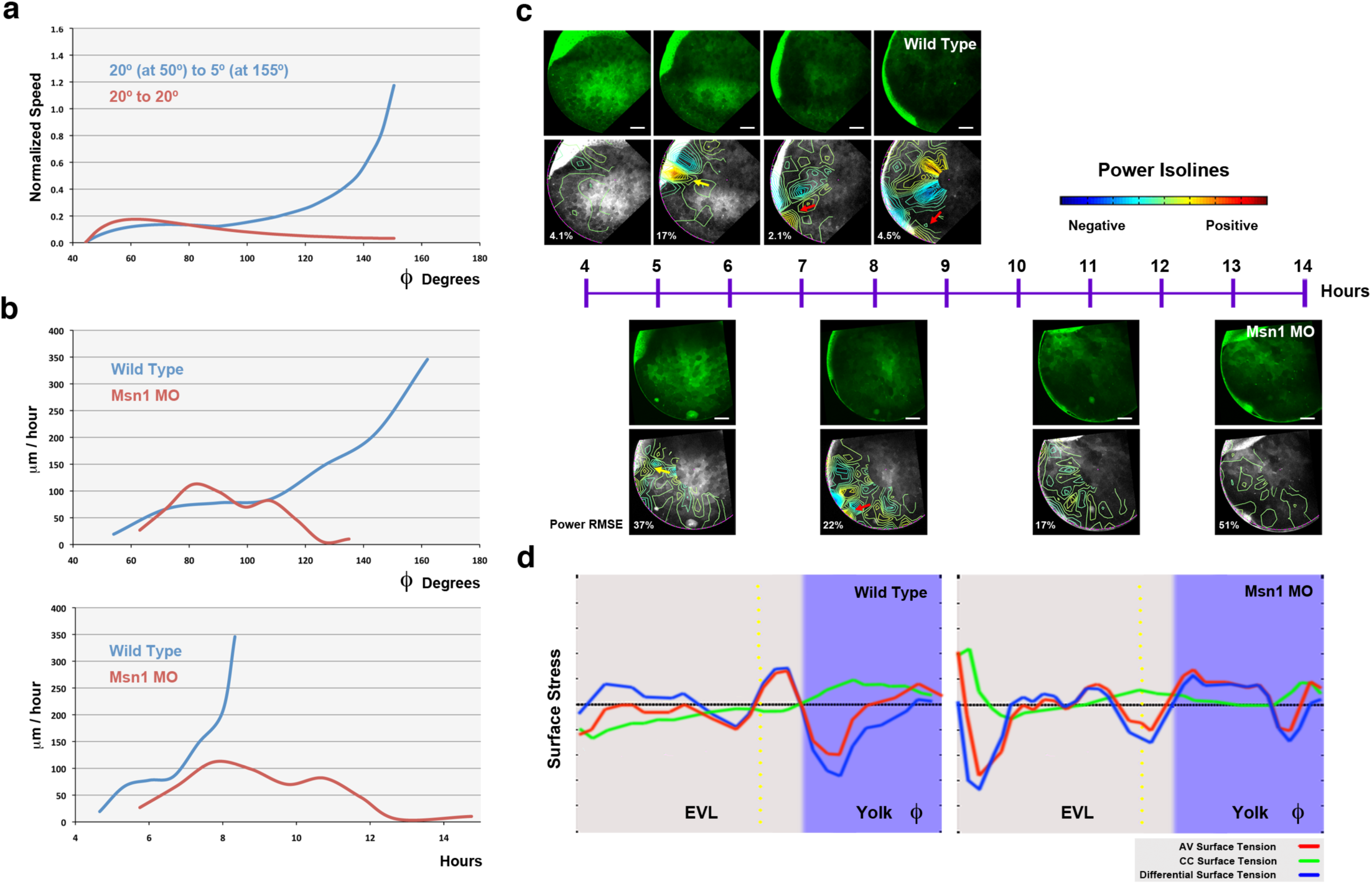
Experimental validation of the differential surface tension model. **(a)** Speed of the EVL advancing front at different epiboly stages (ϕ angle) for two distinct simulated conditions: 1) a difference of elastic tension between the EVL and the yolk and a two-phase reduction of the contractile E-YSL width (from 20° to 5°) as epiboly advances (from 50° to 155°) (wild type - blue) and 2) equivalent difference of elastic tension to 1 but with no shrinkage (constant 20°) of the E-YSL (contractileless - red). **(b)** Experimental values for the EVL advancing front speeds at different epiboly stages (ϕ angle - top) and at different times (bottom) for wild type controls (blue) and Msn1 yolk morphants with reduced E-YSL contractility (red). Experimental profiles closely approach to those inferred in the simulations (compare to **(a)**). **(c)** Timeline showing medial sections and power isolines for wild type (top) and Msn1 yolk morphants (bottom) along epiboly. Mechanical parameters were inferred from experimental velocity fields by HR (**Supplementary Video 1**0). The RMSE of power is shown as a percentage at each time point. DCs in the Msn1 yolk morphants undergo ingression on schedule at 50 % epiboly (yellow arrow). Importantly, as in wild type controls, the active work in morphants is detected at the E-YSL (red arrow). This power supply fades away as epiboly slows down and eventually fails. Scale bar 100 μm. **(d)** Tensional profiles inferred by HR for wild type at 80 % epiboly and its temporal equivalent (9 hours after egg laying) for Msn1 yolk morphants. The characteristic profiles of longitudinal and latitudinal stresses and the animal to vegetal stress gradient that are built up during epiboly are never properly developed in morphants.aYolk Granules

We experimentally tested these inferences by genetically interfering in Msn1 expression, a zebrafish ortholog of the *Drosophila* Ste20-like kinase Misshapen. Msn1 is a well-characterized mediator of E-YSL contractility required for the localized recruitment of actin and myosin 2 within the E-YSL ^21^. We directly addressed the role of *msn1* in the E-YSL by injecting morpholinos in the yolk at the sphere stage. This resulted in epiboly's slowness and arrest with 50% penetrance, as previously described. We then performed a kinematic and mechanical analysis of *msn1* yolk morphants and found that the failure on E-YSL contractility was associated to a strong reduction in the speed of progression of the EVL front mostly after 50 % epiboly (**Fig. 7b** and **Supplementary Video 10**). We further found by HR that the magnitude of the power density at the E-YSL was critically compromised and that the stress gradients along the embryo cortex were essentially abolished (**Fig. 7c** and **d**).

In short, simulations and experimental interferences in E-YSL contractility show a remarkable correlation strongly supporting that epiboly kinematics and mechanical behavior can be minimally described by 1) the contractile capabilities and gradual change of dimensions of the E-YSL and 2) the dissimilar elastic properties of the EVL and the yolk surface **(Fig. 6b)**.

## DISCUSSION

We have performed a detailed study to determine the up to now elusive, major force-generating elements driving zebrafish epiboly ^23, 35^. We revealed several significant evidences: 1) both, the yolk surface (this manuscript and ^12^) and the EVL tissue ^36^ display elastic behavior at short time scales. This elastic response is sustained all throughout epiboly; 2) the yolk cell membrane behaves as an extremely soft solid; 3) the yolk content is fairly viscous and incompressible and flows at very low Reynolds number with a negligible inertia and 4) the magnitudes of the cortical and yolk fluid tractions are comparable and of equivalent order. Further, we generated a theoretical approximation to the embryo as a tensile cortex of spherical shape (SC) surrounding a viscous fluid. Applying Hydrodynamic Regression (HR) to this SC model, we fitted a unique velocity and dynamic pressure field to experimental 2D+T time-lapses and from here we built a minimal mechanical model of epiboly.

HR is a non-invasive, novel method able to quantitatively infer dynamic mechanical parameters (mechanical power density and surface/cortical tension) from *in vivo* imaging. As stands today, the method can, in principle, be applied to any deforming tissue and, with minor modifications, be extended to full 3D analysis. Consequently, HR may become instrumental in the quantitative study of cell and tissue mechanics during embryonic development as well as in physiological and pathological processes. An extended discussion on the HR method applicability and restrictions is presented in **Supplementary Note 8**. The ability of accurately inferring and mapping mechanical power density patterns and surface tension profiles *in vivo* and in whole organisms enables the exploration of the genetic and cellular mechanisms that account for the biomechanics of morphogenesis.

In its application to epiboly, HR enabled us to estimate not only mechanical power densities and provide insight in the effective surface tension variations but also to reach a global concerted representation of epiboly in which cell movements can be explained as a response to mechanics. Before surpassing the equator, an active role for the EVL and DCs on epiboly is supported by the mechanical power densities at the blastoderm (see **Fig. 4d)** and the stresses at the embryo surface (see **Fig. 4e)** uncovered by HR. EVL and DCs early activity is instrumental for epiboly progression ^10, 21, 37, 38^. From 50 % onwards, however, we suggest that epiboly becomes driven by the concerted actions of latitudinal (CC) and longitudinal (AV) forces at the contractile E-YSL and the polarized elastic properties of the surface **(Fig. 6a)**. We showed through simulations and experimental interference that differential tensional properties of the yolk surface and the EVL altogether with the contractile capability of the E-YSL are sufficient to explain both epiboly surface and inner movements at least after 50 % epiboly (see model in **Fig. 6b)**. This model is further backed by the documented preferential orientation of EVL cells which suffer anisotropic tension in the AV direction ^21^. It is also in agreement with recent observations pointing to the EVL as a continuous elastic medium in which tensions and elastic deformations occur at appropriate length scales (see ^36^). This last issue is consistent with equivalent observations in *Fundulus* ^39^.

It has been suggested, alternatively, that epiboly may be described as the result of the combined activities of cable-constriction and flow-friction motors ^12^. In this opposing mechanical scenario, tissue deformation would be driven by the ability of the cortical actomyosin ring at the E-YSL to contract and generate active tension, while an additional pulling longitudinal force in the yolk will be generated by a regime of intermediate friction acting against an actomyosin retrograde flow. This cortical-friction motor proposition derives from a theoretical description of YSL actomyosin network dynamics and it has not been experimentally tested. It implies that the actomyosin ring, which is mechanically connected to the EVL on its animal side, should have a free boundary on its vegetal side. This assumption is essential for the model and demands that the surface tension from the contractile ring towards the vegetal pole should decrease towards zero. We have found, however, that laser microsurgery in the E-YSL unequivocally pointed to a positive oriented gradient of AV tension from the EVL / yolk margin towards the vegetal pole **(Fig. 5c)**. Further, our measurements of surface tension (T_c_) by AFM *in vivo* explicitly revealed a non-null surface/cortical tension at the vegetal pole, which increased with time as epiboly progresses **(Fig. 5e)**. Therefore, the vegetal boundary of the E-YSL is not mechanically free, as it was previously claimed ^12^, but linked to the tense yolk membrane. In these circumstances, the tension originated by the constriction of the actomyosin ring (at the E-YSL) will be conveyed by the yolk cell membrane towards the vegetal pole. Hence, for epiboly progression, a flow-friction motor would be, in principle, needless.

In our view, the previous failure to detect tension at the vegetal pole ^12^, while we readily detect a significant surface tension with AFM strongly suggests that the yolk surface/cortical tension is indeed sustained at the pole by the elastic membrane itself and not by the underlying cortex. Remarkably, considering the notable fitting of our simulations, any potential anisotropy in material properties of the yolk membrane seems not to be mechanically relevant for epiboly. In this context, the role of the observed yolk cortical retrograde flows of actin and myosin might be, not to provide a friction force, but to replenish actin and myosin, which undergo rapid turnover at the contractile E-YSL.

In summary, by applying HR we have defined the spatio-temporal evolution of local cortical tensional stresses and power sources during zebrafish epiboly. We also found that the incompressible yolk displayed stereotyped laminar flows and low but measurable mechanical energy dissipation, suggesting a novel morphogenetic mechanism by which internal flows convey a passive transmission of cortical forces leading to tissue deformation. Our simulations and experimental interferences into E-YSL contractility *(msn1* yolk morphants) (**Fig. 7**) further support that the coordinated action of cortical elements (local contractility at the E-YSL and differential tension between the EVL and the yolk surface) and passive viscous flows in the yolk is instrumental in the mechanical implementation of epiboly morphogenetic movements **(Fig. 6b)**. Still, an active role of the EVL, mostly before reaching the equator, and a mechanical input linked to DCs ingression ^37, 38, 10, 21^ may be necessary to refine the process by regulating precise timing and the shape of cell layers twists.

Local tensional stresses and force passive transmission with elastic and viscous components in an incompressible medium provide a simple physical description for the mechanical control of morphogenetic movements. This biological logic appears to be widespread and it applies, for instance, to the folding of the epithelia in the early *Drosophila* embryo during gastrulation, where stresses generated at the surface integrate with the hydrodynamic properties of the interior to transmit force ^20^. We propose that hydrodynamics may be key for the comprehension of the biomechanics of multiple morphogenetic processes

## METHODS

Methods and any associated references are available in the online version of the paper

## ACKNOWLEDGEMENTS

We thank the Confocal Microscopy Unit from IBMB-PCB, the Advanced Digital Microscopy Core Facility from IRB Barcelona, Xavier Esteban and members of the EMB laboratory for continuous support. We are grateful to Javier Buceta, Carolina Minguilløn, Emmanuel Farge, Damian Brunner, Katerina Karkali and Carla Prat for reading earlier versions of this manuscript and John P. Trinkaus for inspiring work. The Consolidated Groups Program of the Generalitat de Catalunya and DGI and Consolider Grants from the Ministry of Economy and Competitivity of Spain to EMB and DN supported this work.

## AUTHOR CONTRIBUTION

AHV and MM performed all biological tests; PAP designed the modeling and the regression analysis; JC, AHV, MM and PAP performed the laser cuts and data refinement; TL and DN provided the AFM and particle tracking rheology data; ST and IP overview the hydrodynamic analysis and code implementation and EMB designed the study, analyzed the data and wrote the paper. AHV, MM and PAP contributed equally to the study. All authors discussed the results and commented on the manuscript.

## REFERENCES

1. Kiehart, D.P., Galbraith, C.G., Edwards, K.A., Rickoll, W.L. & Montague, R.A. Multiple forces contribute to cell sheet morphogenesis for dorsal closure in Drosophila. J Cell Biol 149, 471–490 (2000).

2. Desprat, N., Supatto, W., Pouille, P.A., Beaurepaire, E. & Farge, E. Tissue deformation modulates twist expression to determine anterior midgut differentiation in Drosophila embryos. Dev Cell 15, 470–477 (2008).

3. Pouille, P.A., Ahmadi, P., Brunet, A.C. & Farge, E. Mechanical signals trigger Myosin II redistribution and mesoderm invagination in Drosophila embryos. Sci Signal 2, ra16 (2009).

4. Rauzi, M. & Lecuit, T. Closing in on mechanisms of tissue morphogenesis. Cell 137, 1183–1185 (2009).

5. Aigouy, B. et al. Cell flow reorients the axis of planar polarity in the wing epithelium of Drosophila. Cell 142, 773–786 (2010).

6. Lepage, S.E. & Bruce, A.E. Zebrafish epiboly: mechanics and mechanisms. The International journal of developmental biology 54, 1213–1228 (2010).

7. Martin, A.C. Pulsation and stabilization: contractile forces that underlie morphogenesis. Developmental biology 341, 114–125 (2010).

8. Grill, S.W. Growing up is stressful: biophysical laws of morphogenesis. Curr Opin Genet Dev 21, 647–652 (2011).

9. Lecuit, T., Lenne, P.F. & Munro, E. Force generation, transmission, and integration during cell and tissue morphogenesis. Annual review of cell and developmental biology 27, 157–184 (2011).

10. Concha, M.L. & Adams, R.J. Oriented cell divisions and cellular morphogenesis in the zebrafish gastrula and neurula: a time-lapse analysis. Development 125, 983–994 (1998).

11. Keller, R., Davidson, L.A. & Shook, D.R. How we are shaped: the biomechanics of gastrulation. Differentiation; research in biological diversity 71, 171–205 (2003).

12. Behrndt, M. et al. Forces driving epithelial spreading in zebrafish gastrulation. Science 338, 257–260 (2012).

13. Kimmel, C.B., Ballard, W.W., Kimmel, S.R., Ullmann, B. & Schilling, T.F. Stages of embryonic development of the zebrafish. Dev Dyn 203, 253–310 (1995).

14. Solnica-Krezel, L. Gastrulation in zebrafish - all just about adhesion? Curr Opin Genet Dev 16, 433–441 (2006).

15. Rohde, L.A. & Heisenberg, C.P. Zebrafish gastrulation: cell movements, signals, and mechanisms. International review of cytology 261, 159–192 (2007).

16. Toyama, Y., Peralta, X.G., Wells, A.R., Kiehart, D.P. & Edwards, G.S. Apoptotic force and tissue dynamics during Drosophila embryogenesis. Science 321, 1683–1686 (2008).

17. Grashoff, C. et al. Measuring mechanical tension across vinculin reveals regulation of focal adhesion dynamics. Nature 466, 263–266 (2010).

18. Rodriguez-Diaz, A. et al. Actomyosin purse strings: renewable resources that make morphogenesis robust and resilient. HFSP J 2, 220–237 (2008).

19. Niwayama, R., Shinohara, K. & Kimura, A. Hydrodynamic property of the cytoplasm is sufficient to mediate cytoplasmic streaming in the Caenorhabditis elegans embryo. Proceedings of the National Academy of Sciences of the United States of America 108, 11900–11905 (2011).

20. He, B., Doubrovinski, K., Polyakov, O. & Wieschaus, E. Apical constriction drives tissue-scale hydrodynamic flow to mediate cell elongation. Nature (2014).

21. Koppen, M., Fernandez, B.G., Carvalho, L., Jacinto, A. & Heisenberg, C.P. Coordinated cell-shape changes control epithelial movement in zebrafish and Drosophila. Development 133, 2671–2681 (2006).

22. Betchaku, T. & Trinkaus, J.P. Contact relations, surface activity, and cortical microfilaments of marginal cells of the enveloping layer and of the yolk syncytial and yolk cytoplasmic layers of fundulus before and during epiboly. J Exp Zool 206, 381–426 (1978).

23. Cheng, J.C., Miller, A.L. & Webb, S.E. Organization and function of microfilaments during late epiboly in zebrafish embryos. Dev Dyn 231, 313–323 (2004).

24. Popgeorgiev, N. et al. The apoptotic regulator Nrz controls cytoskeletal dynamics via the regulation of Ca2+ trafficking in the zebrafish blastula. Dev Cell 20, 663–676 (2011).

25. Supatto, W. et al. In vivo modulation of morphogenetic movements in Drosophila embryos with femtosecond laser pulses. Proceedings of the National Academy of Sciences of the United States of America 102, 1047–1052 (2005).

26. Cartwright, J.H., Piro, O. & Tuval, I. Fluid dynamics in developmental biology: moving fluids that shape ontogeny. HFSP J 3, 77–93 (2009).

27. Daniels, B.R., Masi, B.C. & Wirtz, D. Probing single-cell micromechanics in vivo: the microrheology of C. elegans developing embryos. Biophys J 90, 4712–4719 (2006).

28. Discher, D.E., Mooney, D.J. & Zandstra, P.W. Growth factors, matrices, and forces combine and control stem cells. Science 324, 1673–1677 (2009).

29. Butcher, D.T., Alliston, T. & Weaver, V.M. A tense situation: forcing tumour progression. Nat Rev Cancer 9, 108–122 (2009).

30. Colombelli, J. et al. Mechanosensing in actin stress fibers revealed by a close correlation between force and protein localization. J Cell Sci 122, 1665–1679 (2009).

31. Evans, E. & Yeung, A. Apparent viscosity and cortical tension of blood granulocytes determined by micropipet aspiration. Biophys J 56, 151–160 (1989).

32. Lomakina, E.B., Spillmann, C.M., King, M.R. & Waugh, R.E. Rheological analysis and measurement of neutrophil indentation. Biophysical journal 87, 4246–4258 (2004).

33. Krieg, M. et al. Tensile forces govern germ-layer organization in zebrafish. Nature cell biology 10, 429–436 (2008).

34. Fujimura, Y., Inoue, M., Kondoh, H. & Kinoshita, S. Measurement of Micro-Elasticity within a Fertilized Egg by Using Brillouin Scattering Spectroscopy. Journal of Korean Physical Society 51, 854 (2007).

35. Solnica-Krezel, L. & Driever, W. Microtubule arrays of the zebrafish yolk cell: organization and function during epiboly. Development 120, 2443–2455 (1994).

36. Campinho, P. et al. Tension-oriented cell divisions limit anisotropic tissue tension in epithelial spreading during zebrafish epiboly. Nat Cell Biol 15, 1405–1414 (2013).

37. Trinkaus, J.P. The cellular basis of Fundulus epiboly. Adhesivity of blastula and gastrula cells in culture. Developmental biology 7, 513–532 (1963).

38. Warga, R.M. & Kimmel, C.B. Cell movements during epiboly and gastrulation in zebrafish. Development 108, 569–580 (1990).

39. Weliky, M. & Oster, G. The mechanical basis of cell rearrangement. I. Epithelial morphogenesis during Fundulus epiboly. Development 109, 373–386 (1990).

40. Wirtz, D. Particle-tracking microrheology of living cells: principles and applications. Annual review of biophysics 38, 301–326 (2009).

